# Primordial cardiomyocytes orchestrate myocardial morphogenesis and vascularization but are dispensable for regeneration

**DOI:** 10.64898/2025.12.29.696817

**Authors:** Jisheng Sun, Lu Chen, Jinhu Wang

## Abstract

The vertebrate heart is composed of heterogeneous cardiomyocyte (CM) populations, however, the roles of distinct CM subpopulations in heart development and repair remain poorly defined. Here, using single-cell RNA sequencing analysis of adult zebrafish heart, we identified a unique CM subpopulation marked by the expression of *phlda2*, which is associated with anaerobic metabolism and different from mature CMs which are enriched for oxidative phosphorylation genes. We demonstrated that *phlda2*^+^ cells constituted a primordial CM compartment localized between compact and trabecular muscles. Genetic ablation of *phlda2*^+^ CMs during development severely disrupted heart morphogenesis, leading to defective myocardial trabeculation and compaction, and impaired coronary vascularization. Surprisingly, despite their essential roles in development, the depletion of *phlda2*^+^ CMs didn’t impair myocardial restoration and revascularization following ventricular resection. We found that this was probably due to the limited regenerative capacity of the primordial CMs themselves, as they failed to regenerate after either surgical amputation or genetic ablation. Our findings identify primordial CMs as an organizer for heart morphogenesis but not essential for regeneration, revealing a fundamental difference between developmental and regenerative programs in the vertebrate heart.

## METHODS

### Zebrafish and heart injuries

4-10-month-old outbred EK or EK/AB zebrafish were used for ventricular resection surgeries. To deplete *phlda2*-expressing cells, *phlda2:mCherry-NTR* transgenic zebrafish were used at the age of 6-8 weeks (juvenile) and 4-12 months (adult). Animal density was maintained at ∼4 fish/L in all experiments. The *phlda2:mCherry-NTR* animals and control wild type siblings were treated with vehicle or 10 mM metronidazole (Mtz) (MilliporeSigma) in a 1.5 L mating tank for 12 hours/day to ablate *phlda2*^+^ cells. Transgenic strains described elsewhere include: *gata4:EGFP*^1^*; cmlc2:EGFP*^2^*; tcf21:mCherry*^3^; and *deltaC:EGFP*^4^. All transgenic strains were analyzed as hemizygotes. All animal procedures were performed in accordance with Emory University guidelines.

### Constructs, zebrafish transgenes

*<u>phlda2:EGFP</u>*. The *phlda2* translational start codon in the BAC clone CH211-193F12 was replaced with *EGFP* by Red/ET recombineering technology (GeneBridges). 5′ and 3′ homologous arms flanking the *EGFP* cassette for recombination were a 248-base-pair (bp) fragment upstream of the start codon and a 364-bp fragment downstream, respectively. BACs for transgenesis were purified with a midi-prepare 100 kit (Qiagen) and co-injected with PI-SceI into one-cell-stage zebrafish embryos. The full name of this transgenic line is Tg (*phlda2:EGFP*)*^em21^*.

*<u>phlda2:mCherry-NTR</u>*. The *phlda2* translational start codon in the BAC clone CH211-193F12 was replaced with *mCherry-NTR* by Red/ET recombineering. 5′ and 3′ homologous arms flanking the *mCherry-NTR* cassette for recombination were a 248-bp fragment upstream of the start codon, and a 364-bp fragment downstream, respectively. The full name of this transgenic line is Tg (*phlda2:mCherry-NTR*)*^em22^*.

### Mtz treatment

For conditional ablation of *phlda2*⁺ cardiomyocytes, zebrafish expressing *phlda2:mCherry-NTR* were treated with 10 mM Mtz for 12 hours per day for three consecutive consecutive days. Fish were washed out and maintained in fresh system water after each daily treatment. For developmental analyses, juvenile zebrafish (7–8 weeks post-fertilization,wpf) were subjected to Mtz treatment as described above. Following completion of Mtz exposure, fish were maintained under standard conditions. Hearts were collected at 10 days post-treatment for assessment of gata4 activation, and at 30 days post-treatment for analysis of myocardial structure and coronary vessel development. For regeneration experiments, adult zebrafish (4–6 months old) were similarly treated with Mtz for three consecutive days with daily washout. Three days after the final Mtz treatment, ventricular apex resection was performed. Hearts were harvested at 7 days post-amputation (dpa) for analysis of gata4 activation and early regenerative responses, and at 30 dpa for evaluation of myocardial regeneration and coronary vessel revascularization.

### Histological methods and quantification

Histological analyses were performed on 10 µm cryosections or whole-mount paraformaldehyde-fixed hearts. Sectioned ventricular tissues were imaged using a Zeiss LSM800 confocal scanning microscope with ZEN V3.7. F.

To quantify trabecular muscle loss, three medial longitudinal sections were selected from each heart and imaged. Single optical slices of the ventricle were acquired using a 20x objective lens (1024 × 1024 pixels) at the same ventricular position in sections from *cmlc2:EGFP* hearts. Trabecular muscle loss was quantified as the ratio of *cmlc2*:EGFP⁺ cell area to the total heart area within a region extending 100 μm from the ventricular wall. Quantification of *gata4*:EGFP^+^ expression in adult fish was performed as described^5^.

To quantify coronary expansion in developing hearts from *deltaC:EGFP* fish, whole-mounted specimens were selected, and ventricle images were captured using a 20x objective lens (1,024 x 1,024 pixels). The coronary vessel lengths were measured by ImageJ software. To quantify the vessel broken, the total number of vessel fragments were counted. To quantify coronary vessel expansion in regenerating hearts from *deltaC:EGFP* fish, images of ventricles were captured using a 20x objective lens (1,024 x 1,024 pixels). The GFP signals were measured in pixels by ImageJ software for signals in the injury site.

To quantify *gata4*:EGFP^+^ expression in developing hearts from *gata4:EGFP* fish, whole-mounted specimens were selected and ventricle images were captured using a 20x objective lens (1,024 x 1,024 pixels). The GFP signals were measured in pixels and mean intensity by ImageJ software in the injury site. To quantify *gata4*:EGFP^+^ expression in regenerating hearts from *gata4:EGFP* fish, images of ventricles were captured using a 20x objective lens (1,024 x 1,024 pixels). The GFP signals were measured in pixels by ImageJ software for signals in the injury site.

### AFOG staining

A triple staining with Aniline blue, acid Fuchsin and Orange-G (AFOG) was performed as previously described^6^. Briefly, cryosections were fixed in Bouin’s solution (Sigma Aldrich) for 2 h at 60°C, and one hour at room temperature. The slides were washed for 30 min in tap water and then treated with 1% phosphomolybdic acid (Sigma Aldrich) and 0.25% phosphotungstic acid solution (Sigma Aldrich) for 5 min. They were rinsed with distilled water, and incubated for 5 min with AFOG solution containing 3 g acid Fuchsin (Sigma Aldrich), 2 g Orange G (Sigma Aldrich), 1 g Anilin Blue (Sigma Aldrich) in distilled water, with pH adjusted to 1.1. After 5-min wash with distilled water, the sections were were dehydrated in graded series of ethanol and two changes of xylene, then mounted with Permount mounting medium (Fisher Chemical). Degree of regeneration was scored in a blind-test assessment of AFOG stained sections at 30 dpa. Three sections with evident injury were selected per heart. The criteria for scoring were based on the amount of persisting collagen (blue staining) and the wound closure with a new myocardium. The hearts without persisting collagenous scar were classified as fully regenerated hearts. Hearts with scar remnants that were covered by a new myocardium was considered as incomplete regeneration. The absence of a new myocardium around the wound was considered as a blocked regeneration.

### Single-cell RNA-sequencing

To prepare *cmlc2*^+^ cells for single-cell RNA-sequencing analyses, *cmlc2:EGFP* transgenic fish were raised to adult stages. Adult hearts were collected at four months old. The heart samples were digested with 0.26 U/mL Liberase^TM^ Thermolysin Medium (TM) based on a previously published protocol^7^. Dissociated cells were spun down and live EGFP^+^ cells were sorted by flow cytometry. Isolated cells were sent to Emory Integrated Genomics Core (EIGC) center for 10x single-cell RNA-sequencing. Single-cell RNA-seq libraries were prepared using the Chromium Single Cell 3’ Library & Cell Bead Kit v3.1 (Cat. No. 1000128, 1000127, 120262; 10x Genomics) according to the manufacturer’s protocol. Libraries were sequenced with an Illumina NextSeq550 using mid-output 150-cycle kits according to manufacturer specifications. The newly generated scRNA-seq data were demultiplexed, aligned, and quantified using Cell Ranger Single-Cell Software. The newly generated scRNA-seq data yielded 136,174,297 total reads with an average sequencing depth of 36,168 reads per cell. Preliminary filtered data generated from Cell Ranger were used for downstream analysis by the Seurat R package according to standard workflow.

The scRNA-seq dataset has been submitted to Gene Expression Omnibus (GEO). To review, please use the GEO accession number GSE312374 with the token gbulwsiivdqlven.

## STATISTICAL ANALYSIS

All data are presented as mean ± standard error of the mean (SEM). All statistical analyses were performed using GraphPad Prism 7 software. The Mann-Whitney Rank-Sum test was used for assessing statistical differences between the 2 groups. Results with *p* values < 0.05 were considered statistically significant. Statistical details of experiments can be found in the figures and figure legends.

### Introduction

The vertebrate myocardium is not a homogeneous tissue but consists of cardiomyocyte (CM) subpopulations that are functionally and molecularly different^8,9^. Recent single-cell transcriptomic studies have revealed this cellular heterogeneity in both mammalian and zebrafish hearts, showing clusters of CMs with specialized metabolic, structural, and contractile properties ^10–14^. Although these studies provide the evidence of cellular diversity of the myocardium, the functional significance of these CM subpopulations during heart development and disease remains a central question in cardiac biology.

Humans, like all mammals, lack natural ability to efficiently replace lost myocardium with new contractile tissue, a deficiency that results in the leading cause of morbidity and mortality^15–17^. However, zebrafish possess a remarkable capacity to effectively regenerate the lost CMs after injury^2,6,18–21^, facilitated by non-myocardial cells including epicardial cells^3,5^, immune cells^22,23^, endothelial cells^24–26^ and nerve cells^27^, making zebrafish an excellent model for exploring natural heart regeneration. Previous studies have shown that spared CMs dedifferentiate and proliferate to replace the lost myocardium^2,16,28^. However, it remains unclear whether CM subpopulations contribute equally to myocardial restoration. Furthermore, although the regeneration process has been generally accepted to recapitulate the developmental program^29^, it is unknown whether all CM subpopulations play similar roles in heart morphogenesis and regeneration. Therefore, identifying and characterizing the roles of CM subpopulations are critical for understanding the mechanisms of heart development and repair.

Adult zebrafish myocardium is composed of three main types of CMs with different spatial arrangements: the inner mass of trabecular CMs, the outer layer of compact CMs, and a single-cell-thick layer of primordial CMs that located beneath the compact layer^30–32^. While the trabecular and compact muscles have been extensively studied^30,33–36^, the function of primordial CMs remains elusive due to the lack of genetic tools for their manipulation. Primordial CMs are thought to retain immature characteristics^37^, but their molecular identity and functional contributions to myocardial morphogenesis and regeneration have not been fully elucidated. Here, we performed single-cell RNA sequencing (scRNA-seq) analysis of adult zebrafish heart to define CM subpopulations, and investigated the contribution of primordial CMs to myocardial morphogenesis and regeneration. Our studies reveals that primordial CMs are an essential organizer for myocardial morphogenesis and coronary vascularization, but not required for myocardial restoration and revascularization, highlighting regeneration goes beyond the reactivation of developmental programs.

### Results

#### 1. Characterization of cardiomyocyte heterogeneity in adult zebrafish heart

To characterize the cellular heterogeneity of adult zebrafish myocardium, we performed scRNA-seq with isolated CMs. The cells activating the regulatory sequences of the CM maker *cmlc2* were purified from the hearts of adult cmlc2*:EGFP* animals^2^. We applied stringent quality filtering and discarded a small number of *EGFP*^−^/*cmlc2*^−^ cells, and obtained high-quality transcriptomes of 1668 CMs. The expression of cmlc2 transcripts and EGFP reporter signal in the single-cell dataset further confirmed the enrichment of cardiomyocytes (Fig. S1A and S1B). Unsupervised clustering of these cells identified 3 different clusters (Fig.1A), each with a distinct gene expression patterns. Cluster 1 and 2 showed an enrichment of genes associated with mature CMs, such as *ckmt2a* (Fig.1B) and *ckmt2b* (Fig.S1C)^37^. However, due to their significant overlap, it remains unclear whether these two clusters represent distinct cellular identities of CMs. In contrast, Cluster 3 was marked by prominent expression of *phlda2* (Fig.1C), as well as *actn1* (Fig.1D) indicative of an immature cellular state^37^, and *notch3* (Fig.S1D), reflecting elevated Notch signaling activity^34^. To further validate their spatial distribution, analysis of previously published spatial transcriptomic data^14^ revealed that *phlda2*, *actn1*, and *notch3* were predominantly expressed in the outer region of the heart (Fig.S2A-S2C). To gain insights into the biological characteristics of these cluster, we performed Gene Ontology (GO) enrichment analysis with specifically enriched transcripts in each CM cluster (Fig.1E). Clusters 1 and 2 were enriched for GO terms related to aerobic respiration and oxidative phosphorylation, suggesting the mature CMs which efficiently produce the energy to sustain the constant contraction^38,39^. Notably, Cluster 2 showed additional enrichment for pathways involved in mitochondrial respiratory chain assembly, ATP synthesis, TCA cycle, and ribosome biogenesis, suggesting a relatively higher metabolic and biosynthetic activity state. In contrast, Cluster 1 was enriched for GO terms associated with mitochondrial stress responses, protein degradation, and cytoprotective pathways, indicating a stress-adapted cardiomyocyte state. Cluster 3 showed reduced enrichment of metabolic pathways, consistent with an immature metabolic profile, and was further enriched for genes involved in muscle development and epithelial morphogenesis, suggesting a role in cardiac morphogenesis and tissue organization.

**Figure 1.**
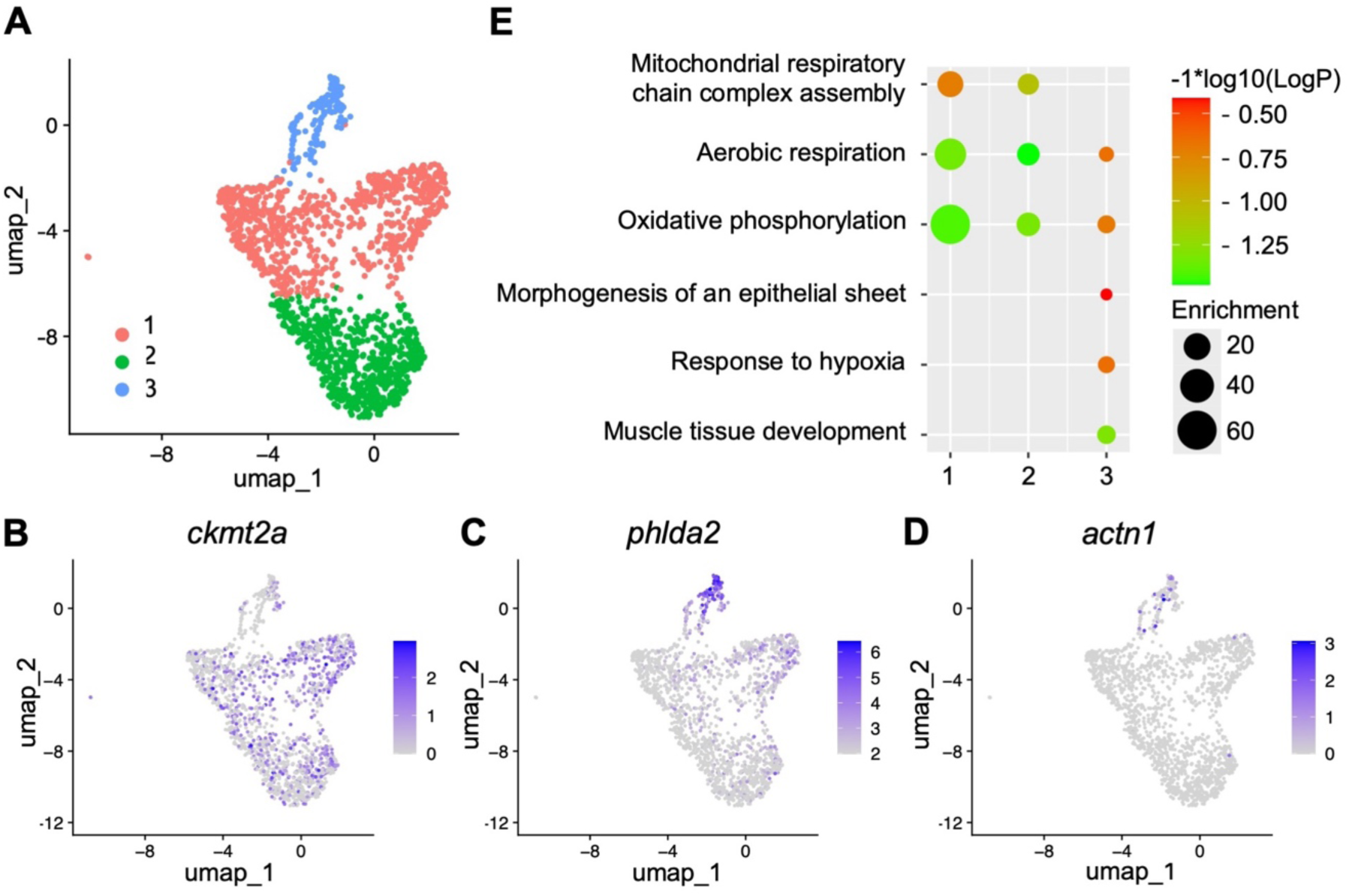
scRNA-seq experiments reveal distinct cardiomyocyte clusters in adult zebrafish hearts. **(A)** Uniform manifold approximation and projection (UMAP) clustering of *cmlc2*^+^ single-cells from adult hearts. **(B-D)** Feature plot of *ckmt2a* **(B)**, *phlda2* **(C)**, *actn1* **(D)**, expression in cardiomyocyte clusters of adult hearts. **(E)** Identification of cardiomyocyte clusters based on gene ontology analysis.

#### 2. The *phlda2*:EGFP^+^ cells represent primordial cardiomyocytes in adult zebrafish hearts

Primordial layer is a single-CM thickness, characteristics of the embryonic ventricle, and high notch activity^30,34,37^. To determine whether Cluster 3 is primordial CMs, we generated *phlda2:EGFP* and *phlda2:mCherry-NTR* BAC transgenic animals, containing sequences of 64,684 bp upstream of the *phlda2* translation initiation codon and 51,710 bp downstream of the stop codon. We observed prominent fluorescence signals in the ventricular wall covering the surface of the ventricle (Fig.2A). As predicted by scRNA-seq result, *phlda2*:EGFP^+^ expression occupied a subset of cells marked by *cmlc2:*mCherry^+^ fluorescence in adult hearts (Fig.2B), indicating that *phlda2*^+^ cells are CMs. Moreover, *phlda2*-directed fluorescence signals did not co-localize with reporter transgenes marking coronary vessels or epicardial cells^3,4^ (Fig.2C and 2D). Examination of juvenile hearts revealed similar expression in the primordial CM layer (Fig. S4A), indicating that *phlda2* transgenic reporters label the primordial cardiac wall throughout the development.

**Figure 2.**
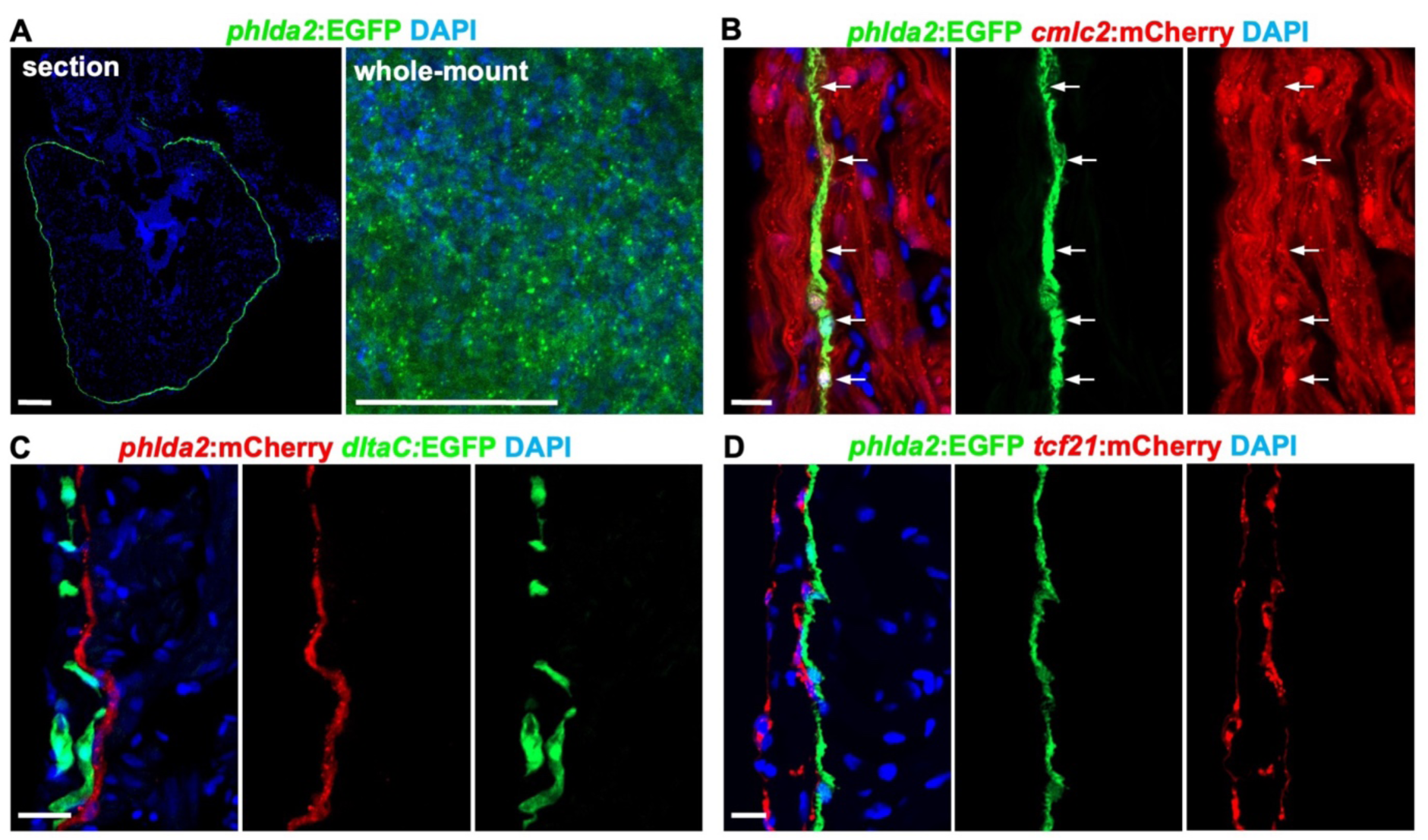
*phlda2*:EGFP^+^ cells specifically label primordial cardiomyocytes in adult zebrafish hearts. **(A)** *phlda2*^+^ cells visualized in whole cardiac sections and whole-mount heart from adult *phlda2:EGFP* animals. n=8. Scale bar, 100 µm. **(B)** Confocal slices indicating *phlda2*^+^ cells in uninjured *phlda2:EGFP;cmlc2:mCherry* ventricles. White arrows represent *phlda2*^+^/*cmlc2*^+^ cells. n=8. Scale bar, 10 µm. **(C)** Confocal slices indicating *phlda2*^+^ cells in adult *phlda2:mCherry;deltaC:EGFP* ventricles. n=12. Scale bars, 20 µm. **(D)** Confocal slices indicating *phlda2*^+^ cells in adult *phlda2:EGFP;tcf21:mCherry* ventricles. n=10. Scale bars, 10 µm. All data are representative of two independent experiments.

#### 3. Primordial cardiomyocytes are required for morphogenesis of myocardium and coronary vascularization

The zebrafish myocardium consists of an inner trabecular layer, a thin primordial layer, and an outer compact layer. Previous multiclonal studies revealed that primordial cardiomyocytes can give rise to trabecular CMs, which later breach the primordial layer to contribute to the compact myocardium^30^. It was reported that nearly 60% of CMs within both the trabecular and outer myocardium originated from primordial cells labeled at the embryonic stage and analyzed at 21 dpf^40^. However, these observations only demonstrate a potential lineage relationship. Whether this cell population is required for the formation or structural integrity of these myocardial layers remains unknown. A major barrier to addressing this question has been the lack of genetic tools for the specific ablation of primordial cells. To address this, we used the bacterial NTR system for inducible cell ablation^41^. We generated a new transgenic line and incubated *phlda2:mCherry-NTR* adults with the prodrug Mtz^41^. This treatment depleted approximately 96.9% of ventricular *phlda2*^+^ cells without apparent effects on animal survival (Fig.S4B). To address the role of *phlda2*⁺ cells during heart development, we performed the following experiments using a standardized Mtz treatment protocol (see Methods), with identical treatment conditions applied to 7-8 wpf juvenile zebrafish. We incubated juvenile *phlda2:mCherry-NTR;cmlc2:EGFP* animal and control *cmlc2:EGFP* siblings for 3 days with Mtz, and then assess the hearts for histological analysis at 10 and 30 days after treatment (dpt). We found primordial CM depletion led to structural abnormality in the heart. The compact myocardium was disorganized compared with controls, and trabecular muscle formation was severely impaired, with an approximately 54.7% reduction in trabecular area, predominantly observed in regions adjacent to the ventricular wall (Fig. 3A, 3B and S3A). In juvenile zebrafish ventricles, *gata4*:EGFP^+^ CMs form the initial clones of compact muscle, which then gradually expand and converge with others to eventually encapsulate the ventricle and create a contiguous wall of compact muscle^35^. We analyzed hearts of *phlda2:mCherry-NTR*;*gata4:EGFP* fish and identified the ablation of primordial cells showed loss of sarcomere organization compared with control siblings at 10 dpt (Fig.3C and S3B). Moreover, primordial cell-depleted hearts exhibited a 16.9% reduction in *gata4*:EGFP fluorescence intensity and a 48.9% reduction in the fluorescence area (Fig.3D-3E).

**Figure 3.**
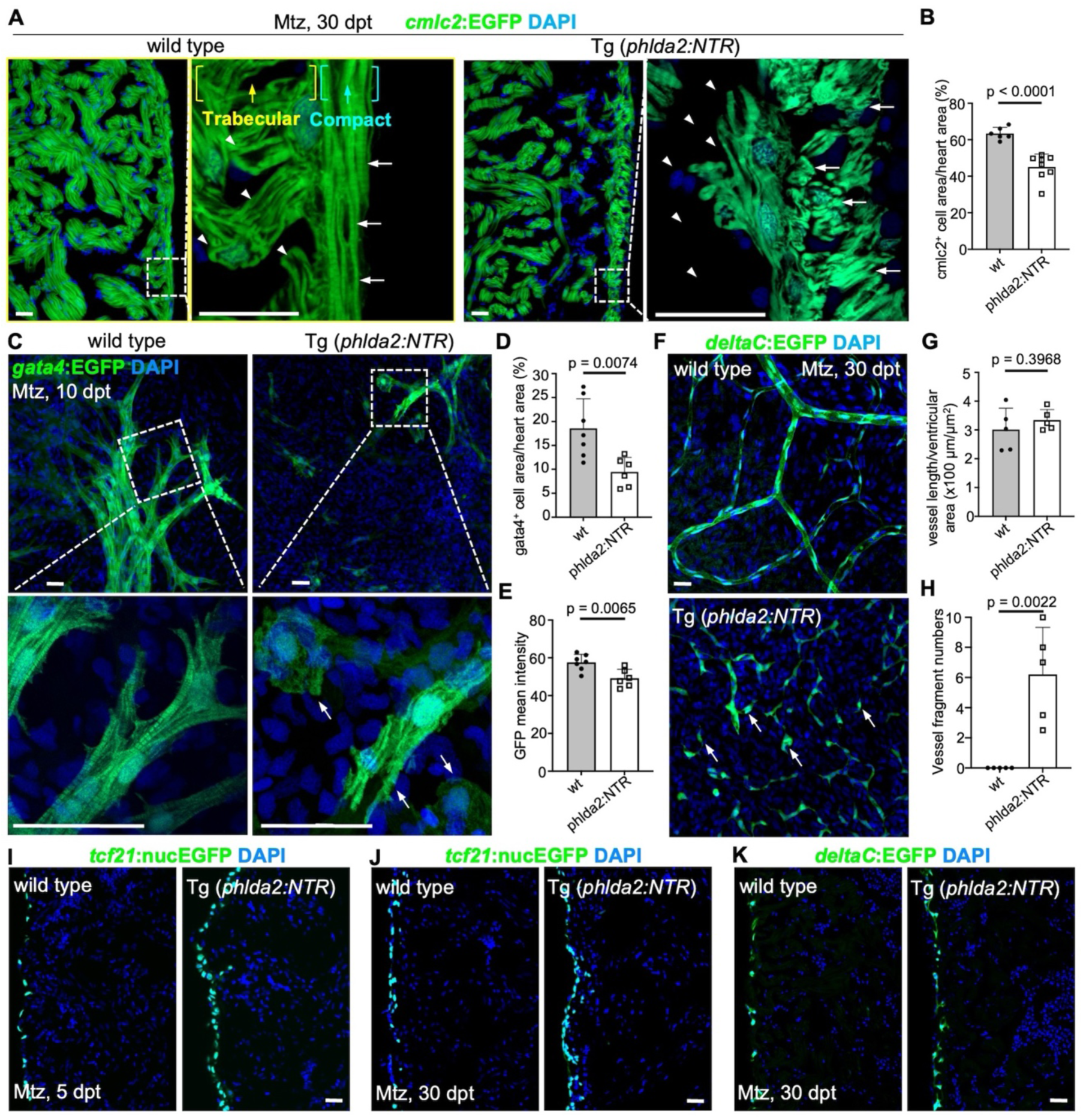
Primordial cardiomyocytes are required for morphogenesis of myocardium and coronary vascularization. **(A)** Compact and trabecular muscle in ventricular sections from Mtz-treated *cmlc2:EGFP* and *cmlc2:EGFP*;*phlda2:mCherry-NTR* animals. Animals were treated with Mtz during the juvenile stage and analyzed 30 dpt. Boxed area is enlarged with high magnification. Arrows represent the compact muscle and arrowheads represent trabecular muscle. N = 6-8. Scale bar, 20 µm. (**B)** Quantification of *cmlc2*⁺ cell area relative to heart area within 100 µm of the heart wall from experiments in (**A)**. Mann-Whitney rank-sum test. **(C)** Visualization of *gata4*^+^ cardiomyocytes in whole-mounted juvenile *gata4:EGFP* and *gata4:EGFP;phlda2:mCherry-NTR* hearts after Mtz treatment. Boxed area is enlarged with high magnification. Arrowheads represent EGFP signals in *phlda2* cell–depleted hearts. n=6-7. Scale bars, 20 µm. (**D-E)** Quantification of the percentage of EGFP^+^ pixels **(D)** and EGFP intensity **(E)** on the ventricular surface from experiments in (**D)** and (**E)**. Mann-Whitney rank-sum test. **(F)** Coronary vessels in whole-mounted *deltaC:EGFP* and *deltaC:EGFP;phlda2:mCherry-NTR* hearts. Animals were treated with Mtz during the juvenile stage and analyzed at 30 dpt. Arrows represent the broken vessel fragments. n=5. Scale bars, 20 µm. **(G-H)** Quantification of vessel length/heart area (**G)** and vessel fragments **(H)** on the ventricular surface from experiments in (**F)**. Vessel length was measured within a fixed region of interest (ROI) after skeletonization of *deltaC:EGFP*⁺ vessels in ImageJ. **(I-J)** Visualization of *tcf21*^+^ epicardial cells in sectioned *tcf21:nucEGFP* and *tcf21:nucEGFP;phlda2:mCherry-NTR* hearts. Animals were treated with Mtz during the juvenile stage and analyzed at 5 and 30 dpt. n=5. Scale bars, 20 µm. **(K)** Coronary vessels in sectioned *deltaC:EGFP* and *deltaC:EGFP;phlda2:mCherry-NTR* hearts. Animals were treated with Mtz during the juvenile stage and analyzed at 30 dpt. n=5. Scale bars, 20 µm. All data are representative of three independent experiments.

During zebrafish heart development, the ventricle becomes vascularized after the compact myocardium has formed post-embryonically^42^. This process is closely associated with myocardial maturation and likely depends on proper organization of the compact layer to provide structural and molecular cues for vessel growth and integrity. To determine whether ablation of primordial CMs affects vascularization during heart development, we examined coronary vessel formation in *phlda2:NTR;deltaC:EGFP* fish, in which *deltaC:EGFP* has been utilized to visualize coronary endothelial cells^4,43^. We found that coronary vessels were fragmented and poorly developed in primordial cell-ablated hearts, although the total vessel length showed no reduction (Fig.3F-3H and S3C). Notably, these vascular abnormalities persisted at 90 days post-treatment, indicating that the defects do not resolve during subsequent cardiac growth and maturation (Fig. S4C). Because epicardial cells play critical roles in coronary vessel development, we next asked whether the vascular defects observed following primordial CM ablation were secondary to alterations in the epicardium. We found that *tcf21*^+^ epicardial cells revealed an increase rather than a decrease in epicardial cell abundance in primordial CM-ablated hearts (Fig. 3I-3J). These findings suggest that the impaired coronary vessel organization is unlikely to result from a reduction in epicardial cell number, although we cannot exclude the possibility that altered epicardial function or signaling contributes to the vascular phenotype. In addition, because primordial cardiomyocytes form a continuous layer at the ventricular surface, we investigated whether their ablation permits abnormal invasion of epicardial cells or coronary vessels into the myocardium. However, we observed no evidence of ectopic localization of either cell population following primordial CM ablation (Fig. 3J-3K). Thus, loss of the primordial CM layer does not appear to disrupt tissue compartmentalization or permit abnormal cellular invasion into the myocardium. Together, these findings demonstrate that primordial CMs are required for proper myocardial growth and coronary vascularization during heart development.

#### 4. Primordial cardiomyocytes are not essential for myocardial regeneration and coronary revascularization

Our data indicated primordial CMs play essential roles during heart development. Next, we examined their contribution to heart regeneration. To address the role of primordial cells during heart regeneration, we perform the below experiments using standardized Mtz treatment protocol (see Methods), with identical treatment conditions applied to adult zebrafish (4–6 months post-fertilization). The *gata4* expression is induced in regenerating CMs and represent the proliferating CMs with reactivation of a cardiac developmental program^2,5,35^. To determine whether ablation of primordial cardiomyocytes is sufficient to activate regenerative signaling, we first treated adult *phlda2:mCherry-NTR;gata4:EGFP* fish and control *gata4:EGFP* siblings with Mtz for 12 hours per day over three consecutive days without ventricular resection. We did not observe any induction of gata4:EGFP expression following primordial CM ablation (Fig. S5), indicating that loss of *phlda2*⁺ cardiomyocytes is not sufficient to trigger regenerative gata4 activation in the absence of injury. Then we treated adult *phlda2:mCherry-NTR;gata4:EGFP* fish and control *gata4:EGFP* siblings with Mtz for 12 hours per day over three consecutive days, starting 6 days before resection of the ventricular apex, and collected hearts at 7 dpa. We imaged and quantified *gata4*:EGFP^+^ signals at the injury site and didn’t detect significant difference in EGFP^+^ signals between *phlda2^+^* cell-depleted fish and control siblings (Fig.4A-4B). In addition, with *cmlc2:EGFP* animals, we found that ablation of primordial CMs did not affect the regeneration index score. Myocardial regeneration proceeded normally, with new muscle effectively replacing the injured tissue by 30 dpa (Fig.4C-4D). To further assess the requirement of primordial CMs for coronary revascularization, we depleted primordial cells in *phlda2:NTR;deltaC:EGFP* fish and didn’t observe found obvious difference of coronary vessel density in the regenerating area of control siblings (Fig.4E-4F). Previous studies have shown that zebrafish hearts can fully regenerate without remaining scars by 30 dpa. We also performed AFOG staining, which labels intact muscle in orange, collagen in blue, and fibrin in red^6,44^. Results showed minimal scar area in primordial cell-depleted hearts, with no obvious difference compared to sibling controls at 30 dpa (Fig.4G-4H). Overall, these findings demonstrate that primordial CMs are not essential for heart regeneration, despite their critical role in development.

**Figure 4.**
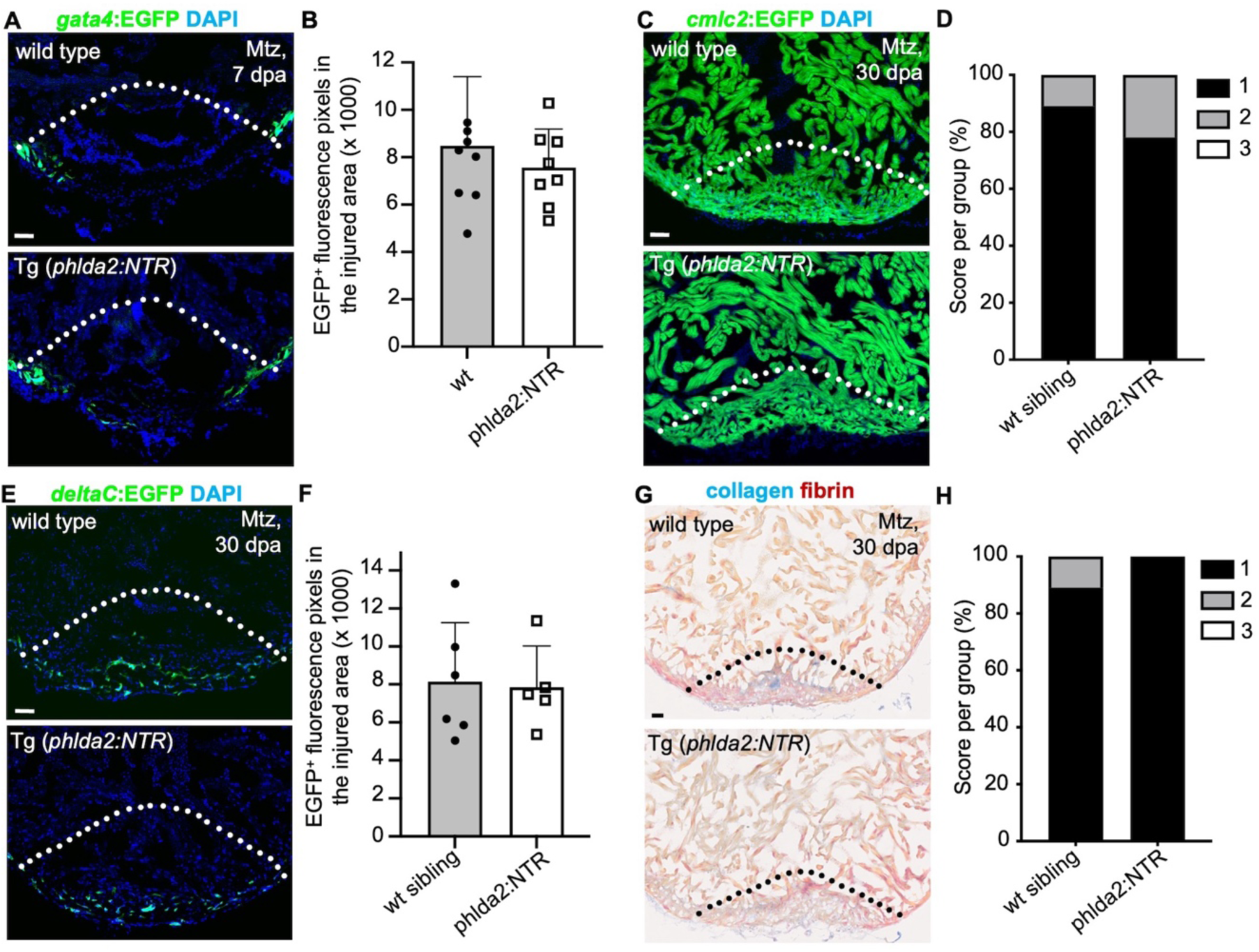
Primordial cardiomyocytes are not essential for myocardial regeneration and coronary revascularization. **(A)** Section images of Mtz-treated *gata4:EGFP* or *gata4:EGFP*;*phlda2:mCherry-NTR* ventricles at 7 dpa assessed for *gata4*:EGFP^+^ CMs in the injury site. Adult animals were treated with Mtz before amputation. Dashed line indicates amputation plane. N = 8. Scale bars, 20 µm. **(B)** Quantification of EGFP^+^ pixels from experiments in **(A).** Scale bars, 20 µm. Mann-Whitney rank-sum test. **(C)**Section images of Mtz-treated *cmlc2:EGFP* or *cmlc2:EGFP*;*phlda2:mCherry-NTR* ventricles at 30 dpa assessed for muscle recovery. Adult animals were treated with Mtz before amputation. Dashed line indicates amputation plane. N = 9 - 10. Scale bar, 20 µm. **(D)** Quantification of regeneration indices from experiments in (**C).** Myocardial regeneration is categorized as follows: 1 = complete regeneration of a new myocardial wall; 2 = partial regeneration; and 3 = a strong block in regeneration. (**E)** Section images of Mtz-treated *deltaC:EGFP* or *deltaC:EGFP*;*phlda2:mCherry-NTR* ventricles at 30 dpa assessed for coronary vessels in the injury site. Adult animals were treated with Mtz before amputation. Dashed line indicates amputation plane. N = 5 - 6. Scale bars, 20 µm. **(F)** Quantification of EGFP^+^ pixels from experiments in **(E).** Scale bars, 20 µm. Mann-Whitney rank-sum test. **(G)** Section images of Mtz-treated *wt siblings* or *phlda2:mCherry-NTR* ventricles at 30 dpa assessed for AFOG staining collagen/fibrin deposition in the injury site. Adult animals were treated with Mtz before amputation. Dashed line indicates amputation plane. N = 9. Scale bars, 20 µm. (H) Quantification of scar score from experiments in **(G).** The scar score is categorized as follows: 1 = hearts without persisting collagenous scar; 2 = hearts with scar remnants that were covered by a new myocardium; 3 = the absence of a new myocardium around the wound. The Chi-squared test was performed. All data are representative of three independent experiments.

#### 5. Primordial cardiomyocytes exhibit limited regenerative capacity

Previous studies have indicated that in adult zebrafish heart, existing CMs can robustly regenerate within several weeks after resection surgery or genetic ablation^18^. We next assessed the regeneration capacity of primordial CMs. We first examined the *phlda2:mCherry-NTR* hearts at various time points after partial resection of the ventricular apex. The primordial layers were lost at the injury site, even 60 dpa at which time the heart regeneration was complete (Fig.5A). We next asked whether this limited regenerative capacity reflects an intrinsic limitation of the cells themselves. To test it, we treat the adult *phlda2:mCherry-NTR* animals without resection with Mtz to ablate primordial CMs. We didn’t observe the recovery of phlda2:mCherry fluorescence, indicating the failure of the primordial regeneration (Fig.5B). Similarly, when juvenile zebrafish (7–8 wpf) were subjected to Mtz-mediated ablation and analyzed 90 days after treatment, *phlda2*^+^ cells remained absent, demonstrating a persistent failure of primordial CM recovery (Fig. S6). Since primordial CMs did not regenerate after either apical resection or direct ablation, we therefore asked whether a regenerative cue could overcome this limitation. We performed apical resection after the ablation of primordial CMs, and found that primordial CMs still failed to regenerate (Fig.5C), suggesting that their limited regenerative capacity is intrinsic rather than environment dependent. Finally, we found that *phlda2*:EGFP^+^ cells in the injury site did not colocalize with *gata4*:EGFP^+^ cells, suggesting they have limited proliferative capacity and do not contribute to the new muscle (Fig. 5D). Consistently, no overlap between *phlda2*⁺ and *gata4*⁺ cardiomyocytes was observed in the 7-8 wpf juvenile zebrafish heart (Fig. 5E). We conclude that the primordial CMs have intrinsically limited regenerative potential, explaining its absence and functional dispensability in the regenerated heart.

**Figure 5.**
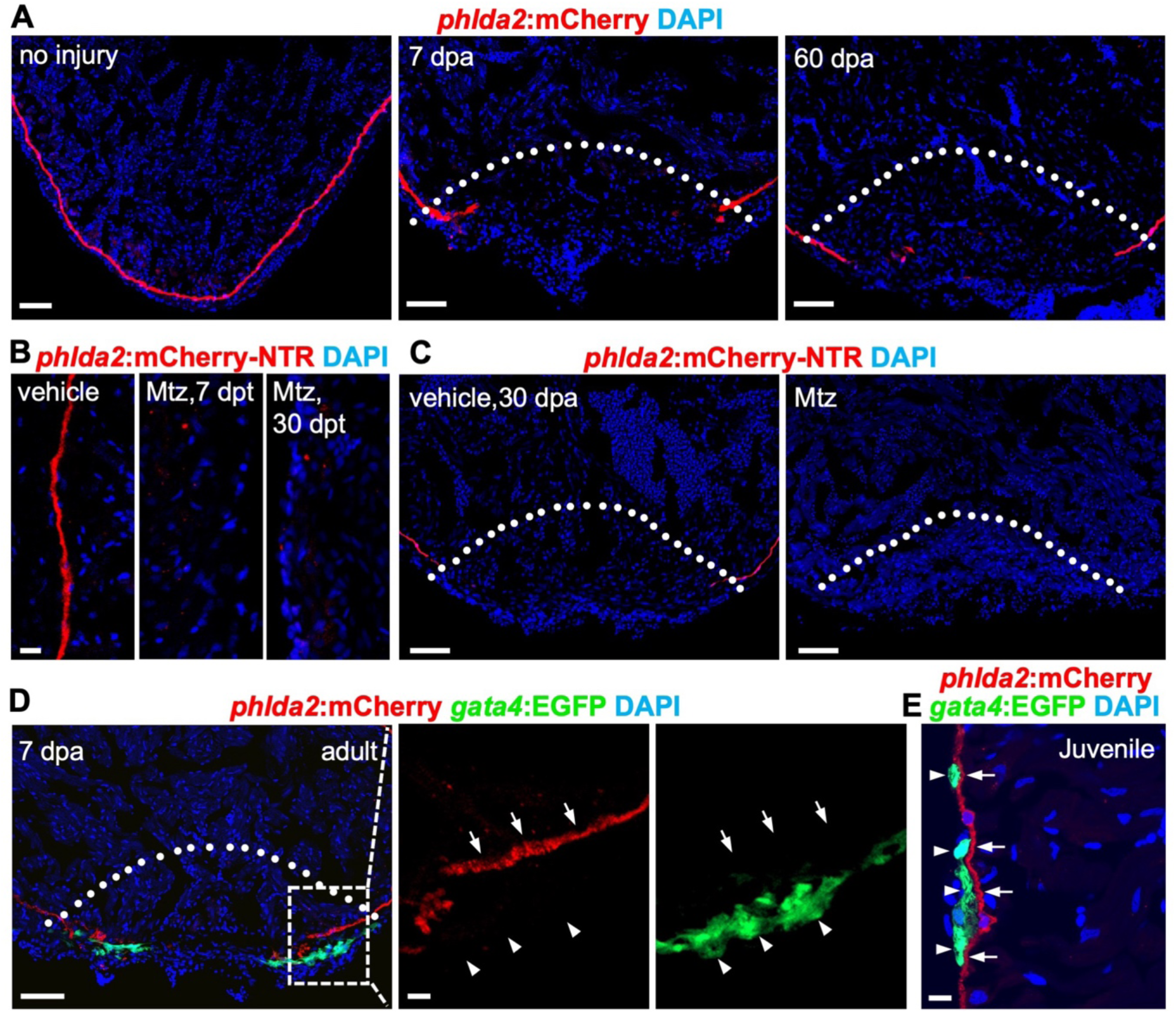
Primordial cardiomyocytes have limited regenerative capacity. **(A)** Section images of *phlda2:mCherry-NTR* ventricles without injury and during regeneration at 7, and 60 dpa. n *=*11-13. Dashed line indicates amputation plane. Scale bar, 20 μm. **(B)** Section images of *phlda2:mCherry-NTR* ventricles without Mtz treatment and after 7 or 30 days of Mtz treatment. n = 12-15. Scale bar, 20 μm. **(C)** Section images of ventricles of vehicle- or Mtz-treated *phlda2a:mCherry-NTR* at 30 dpa. Animals were treated with Mtz during the juvenile stage and amputated as adults. Dashed line indicates amputation plane. n = 12. Scale bars, 20 µm. **(D)** Section images of *phlda2:mCherry-NTR;gata4:EGFP* ventricles at 7 dpa assessed for *phlda2*^+^ cells and *gata4*:EGFP^+^ CMs in the injury site. Box area is shown in higher magnification. Arrows represent *phlda2*^+^ cells; arrowheads represent *gata4*^+^ cells. Dashed line indicates amputation plane. n = 12. Scale bar, 20 µm. (E) Section images of juvenile *phlda2:mCherry-NTR;gata4:EGFP* ventricles assessed for *phlda2*^+^ cells and *gata4*:EGFP^+^ CMs. Arrows represent *phlda2*^+^ cells; arrowheads represent *gata4*^+^ cells. n = 6. Scale bar, 10 µm. All data are representative of two independent experiments.

#### Discussion

In this study, we characterized CM subpopulations in the adult zebrafish heart using scRNA-seq and identified *phlda2* as a marker of primordial CMs, which exhibit transcriptional features of an immature cellular state and functions associated with morphogenesis of an epithelial sheet. Using newly generated *phlda2* reporter and ablation lines, we revealed that primordial CMs reside beneath the compact layer and play essential roles in ventricular morphogenesis and coronary vascularization. However, unlike other CM subtypes, these cells display very limited regenerative capacity and are dispensable for myocardial regeneration. Our findings identified primordial CMs as a crucial organizer for heart morphogenesis but plays no roles during regeneration, revealing a difference between developmental and regenerative programs in the vertebrate heart.

During heart development, multicolor clonal analyses have revealed that the primordial CM lineage contributes to trabecular formation, and trabecular CMs later breach this layer to form the compact myocardium ^30,45,46^. However, direct functional evidence has been lacking due to the absence of specific genetic tools. In this study, we generated a *phlda2:mCherry-NTR* line to specifically deplete primordial CMs. The results show that the depletion of these cells caused a marked reduction in the outer trabecular muscle, whereas inner trabeculae remained largely intact, thereby supporting previous reports that primordial CMs, positioned outside the trabecular layer, contribute to the formation of the trabeculae^30^. Furthermore, we observed a disorganized compact myocardium and disrupted *gata4* expression after the loss of primordial CMs, indicating a crucial role for primordial CMs in coordinating myocardial structural organization and maturation. The loss of these cells likely disrupts the structural and molecular cues required for compact muscle maturation, resulting in an immature ventricular wall.

Coronary vessel formation was also severely affected by the loss of primordial CMs, leading to fragmented and poorly developed vasculature. The coronary vasculature normally invades the compact myocardium during post-embryonic development, a process tightly coupled with myocardial maturation^24,42^. Thus, the vascular defects we observed could arise indirectly from the disorganization of the compact myocardium, which fails to provide a proper scaffold for vessel growth. Alternatively, primordial CMs themselves may produce paracrine factors that promote vessel growth and stability. Consistently, our scRNA-seq analysis revealed that *vegfaa* was highly expressed in primordial CMs (Fig. S1E). VEGF signaling plays a pivotal role in vascular development, acting as an important endothelial mitogen that stimulates endothelial cell tubulogenesis, and maintains vessel stability and integrity^25,47,48^. These findings suggest that primordial CMs may serve as a critical architectural and signaling hub to coordinate the morphogenesis of both myocardial and vascular structures during heart development. Although primordial CM ablation does not affect survival under laboratory conditions, the observed structural defects may have functional consequences on cardiac performance and overall physiological fitness, which warrant further investigation.

In contrast to their essential role in heart development, primordial CMs appear to play no role in cardiac regeneration. Depletion of these cells did not affect myocardial or vascular regeneration, nor did it impair scar resolution. Furthermore, primordial CMs failed to regenerate after either surgical amputation or genetic ablation, even under strong regenerative conditions. Their inability to regenerate in the injured tissue, together with the lack of overlap with *gata4*⁺ proliferative CMs, indicates that these cells possess intrinsically limited proliferative capacity. They are different from trabecular and compact CMs, which efficiently contribute to the formation of new myocardium. This intrinsic restriction likely explains why the loss of primordial CMs does not impair the heart regeneration, suggesting that zebrafish cardiac regeneration is primarily driven by other CM subsets with greater regenerative potential.

Our study demonstrates that primordial CMs are essential for heart morphogenesis but dispensable for regeneration. Although a direct corollary of *phlda2*⁺ primordial cardiomyocytes has not yet been identified in mammals, mammalian hearts contain heterogeneous cardiomyocyte populations with immature states. Whether these populations represent a functional equivalent of zebrafish primordial CMs remains an open question and needs further investigation. These findings emphasize the functional diversity among CM subtypes and broaden our knowledge of CM heterogeneity in the zebrafish heart. Importantly, our results show that heart regeneration does not simply reactivate the developmental programs across all cardiomyocytes. Instead, regeneration depends on the CM subsets with high proliferative and regenerative capacity, rather than engaging all CMs equally. This work provides new insights into heart development and regeneration and may inspire new ideas to enhance cardiac repair in mammals.

## Data availability

All data generated or analyzed during this study are included in the manuscript and supporting files; source data files are available from Dr. Jisheng Sun upon request.

## ACKNOWLEDGMENTS

We thank the Developmental Studies Hybridoma Bank for antibodies. Microscopy data for this study were acquired and analyzed using the Microscopy in Medicine Core in Cardiology at Emory. This work was supported by grants from NHLBI (2R01HL142762) and AHA (25TPA1474053) to J.W. This work was supported by the

NIH National Institute on Aging through the Emory Specialized Center of Research Excellence on Sex Differences (U54AG062334) to J.S. The Microscopy in Medicine Core in Cardiology at Emory was supported by the NIH grant (P01 HL095070).

## AUTHOR CONTRIBUTIONS

Conceptualization, J.S., and J.W.; methodology, J.S., and J.W.; investigation, J.S., L.C., and J.W.; formal analysis, J.S., L.C., and J.W.; visualization, J.S., L.C., and J.W.; writing, J.S. and J.W.; funding acquisition, J.W.; supervision, J.W. All authors commented on the manuscript.

