## Supplemental Figures for "Primordial cardiomyocytes orchestrate myocardial morphogenesis and vascularization but are dispensable for regeneration"

### Figure S1

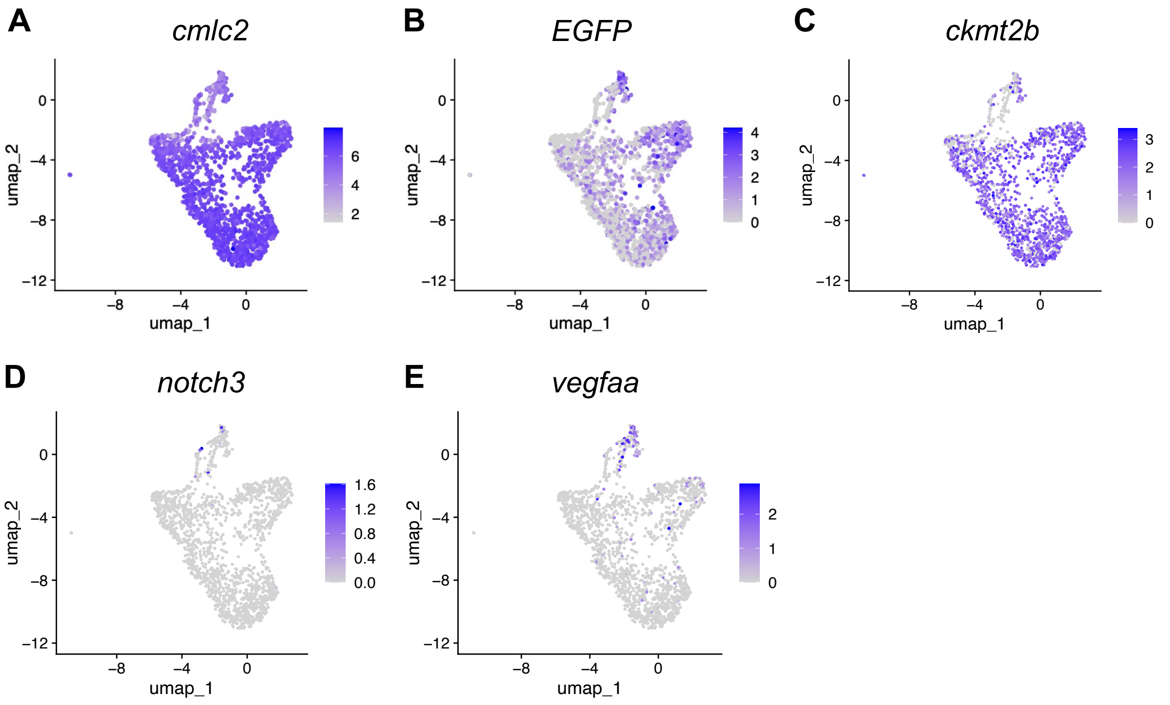


**Figure S1.** Feature plot of *cmlc2* **(A)**, *EGFP* **(B)**, *ckmt2b* **(C)**, *notch3* **(D)**, *vegfaa* **(E)**, expression in cardiomyocyte clusters of adult hearts.

#### Figure S2

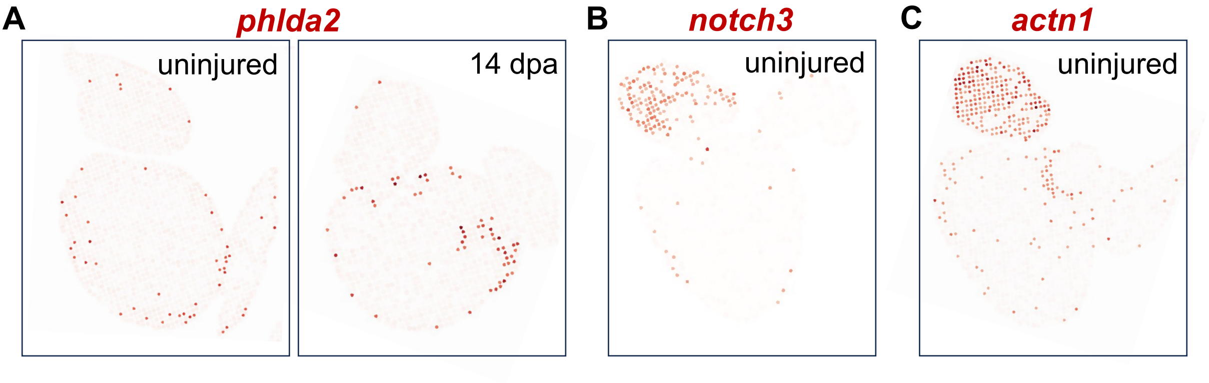


**Figure S2. Spatial visualization of imputed genes using a published zebrafish heart regeneration spatial transcriptomic dataset. (A)** Spatial visualization of imputed expression of *phlda2* in uninjured (left) and regenerated hearts at 17 dpa (right). **(B-C)** Spatial visualization of imputed expression of *notch3* **(B)** and *actn1* **(C)** in uninjured hearts.

#### Figure S3

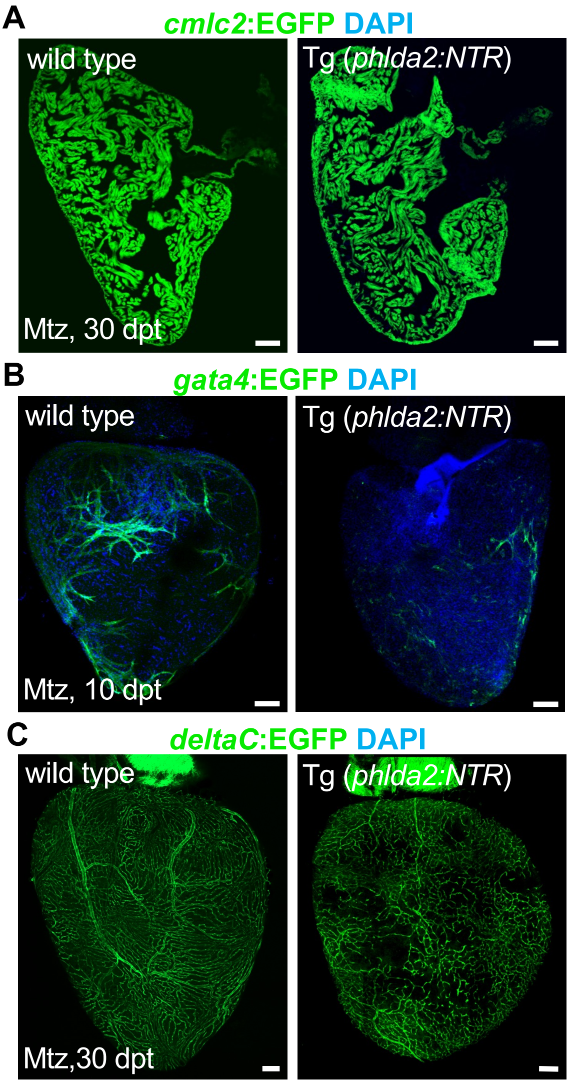


**Figure S3. Low-magnification images showing the entire heart following juvenile-stage ablation of *phlda2*$^{+}$ cardiomyocytes. (A)** Compact and trabecular muscle in ventricular sections from Mtz-treated *cmlc2:EGFP* and *cmlc2:EGFP;phlda2:mCherry-NTR* animals. Animals were treated with Mtz during the juvenile stage and analyzed 30 dpt. Scale bar, 100 µm. **(B)** Visualization of *gata4*$^{+}$ cardiomyocytes in whole-mounted juvenile *gata4:EGFP* and *gata4:EGFP;phlda2:mCherry-NTR* hearts after Mtz treatment.. Scale bars, 100 µm. **(C)** Coronary vessels in whole-mounted *deltaC:EGFP* and *deltaC:EGFP;phlda2:mCherry-NTR* hearts. Animals were treated with Mtz during the juvenile stage and analyzed at 30 dpt. Scale bars, 100 µm. All data are representative of two independent experiments.

### Figure S4

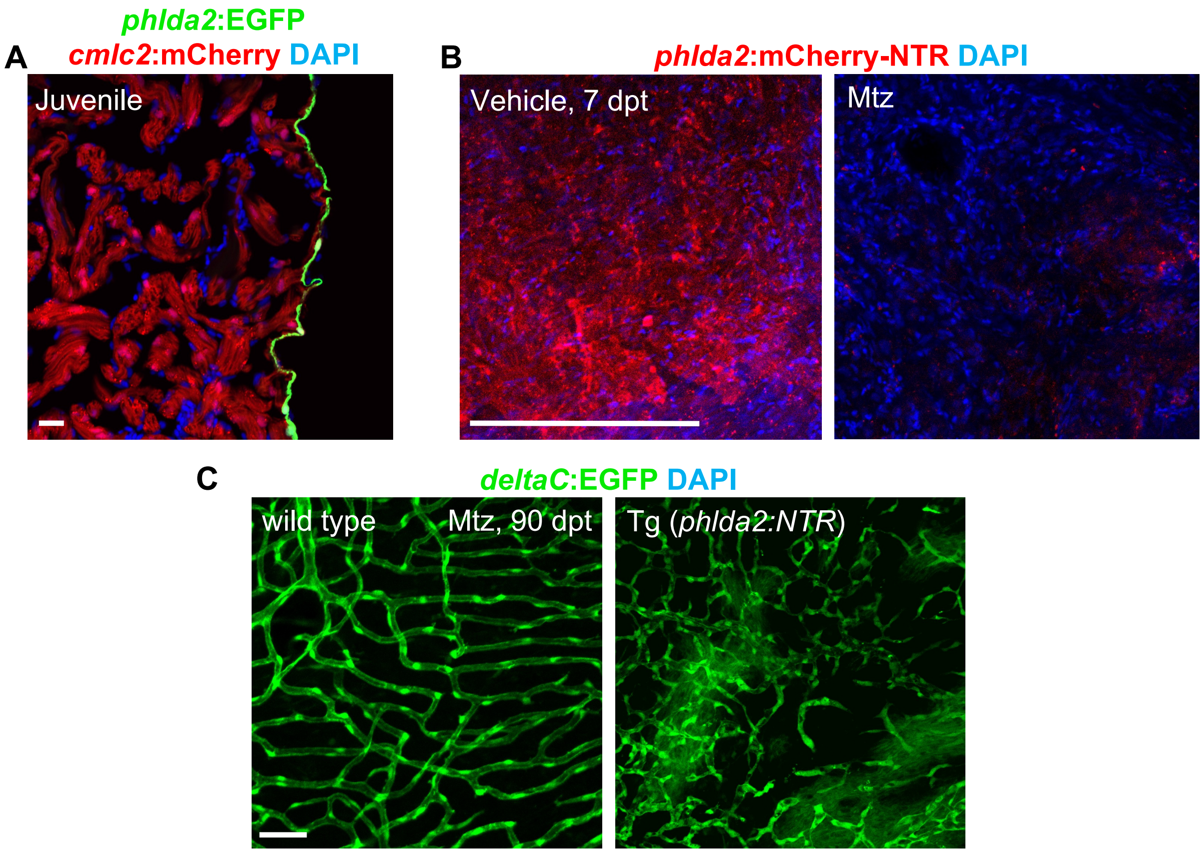


**Figure S4. Validation of *phlda2*⁺ cardiomyocyte ablation in juvenile hearts.** **(A)** Confocal slices indicating *phlda2*⁺ cells in juvenile *phlda2:EGFP; cmlc2:mCherry* ventricles. n = 10. Scale bar, 20 μm. **(B)** Visualization of *phlda2*⁺ cells in whole-mounted adult *phlda2:mCherry-NTR* hearts after vehicle or Mtz treatment. n = 12. Scale bars, 20 μm. **(C)** Coronary vessels in whole-mounted *deltaC:EGFP* and *deltaC:EGFP;phlda2:mCherry-NTR* hearts. Animals were treated with Mtz during the juvenile stage and analyzed at 90 dpt. n=5. Scale bars, 50 μm. All data are representative of two independent experiments.

### Figure S5

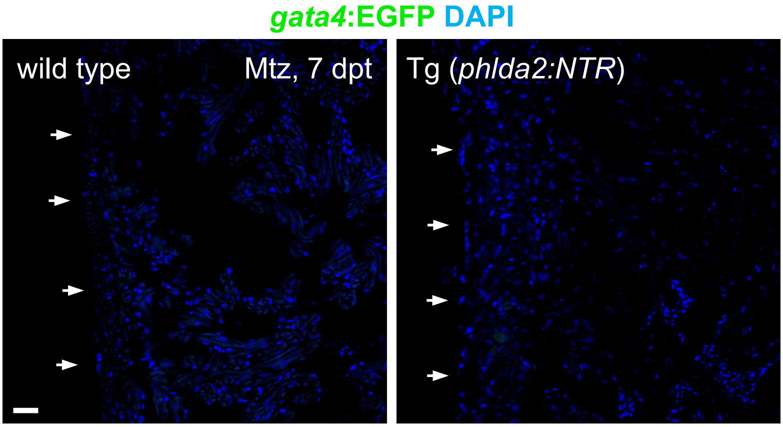


**Figure S5.** Visualization of *gata4*$^{+}$ cardiomyocytes in sectioned adult *gata4:EGFP* and *gata4:EGFP;phlda2:mCherry-NTR* hearts after Mtz treatment. n = 6. Scale bars, 20 μm. All data are representative of two independent experiments.

#### Figure S6

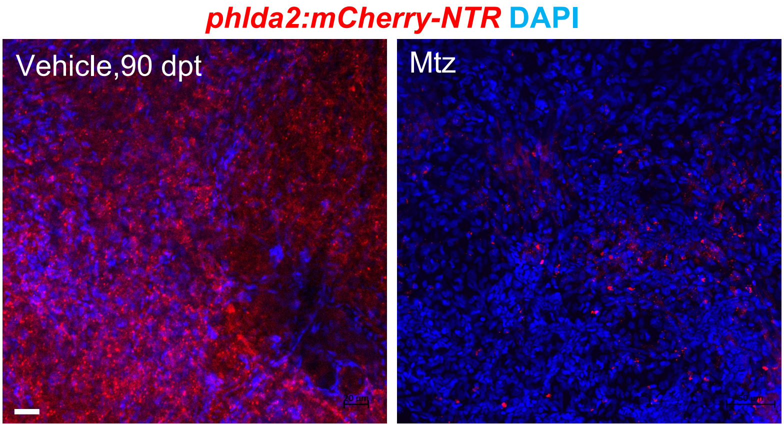


**Figure S6.** Visualization of *phlda2*$^+$ cells in whole-mounted adult *phlda2:mCherry-NTR* hearts after vehicle or Mtz treatment. n = 5. Scale bars, 20 μm. All data are representative of two independent experiments.
